# Web-based Psychoacoustics: Hearing Screening, Infrastructure, and Validation

**DOI:** 10.1101/2021.05.10.443520

**Authors:** Brittany A. Mok, Vibha Viswanathan, Agudemu Borjigin, Ravinderjit Singh, Homeira Kafi, Hari M. Bharadwaj

## Abstract

Anonymous web-based experiments are increasingly and successfully used in many domains of behavioral research. However, online studies of auditory perception, especially of psychoacoustic phenomena pertaining to low-level sensory processing, are challenging because of limited available control of the acoustics, and the unknown hearing status of participants. Here, we outline our approach to mitigate these challenges and validate our procedures by comparing web-based measurements to lab-based data on a range of classic psychoacoustic tasks. Individual tasks were created using jsPsych, an open-source javascript front-end library. Dynamic sequences of psychoacoustic tasks were implemented using Django, an open-source library for web applications, and combined with consent pages, questionnaires, and debriefing pages. Subjects were recruited via Prolific, a web-based human-subject marketplace. Guided by a meta-analysis of normative data, we developed and validated a screening procedure to select participants for (putative) normal-hearing status; this procedure combined thresholding of scores in a suprathreshold cocktail-party task with filtering based on survey responses. Headphone use was standardized by supplementing procedures from prior literature with a binaural hearing task. Individuals meeting all criteria were re-invited to complete a range of classic psychoacoustic tasks. Performance trends observed in re-invited participants were in excellent agreement with lab-based data for fundamental frequency discrimination, gap detection, sensitivity to interaural time delay and level difference, comodulation masking release, word identification, and consonant confusions. Our results suggest that web-based psychoacoustics is a viable complement to lab-based research. Source code for our infrastructure is also provided.

## Introduction

Behavioral experiments that assess the ability of human listeners to detect and discriminate various acoustic cues have provided considerable insight into the psychology of human auditory perception (Moore, 2012), placed useful constraints on theories about the neural coding of sound (Oxenham, 2018), revealed our ability to use acoustic regularities for perceptual organization of sound (Bregman, 1990), and yielded models of speech intelligibility (Bronkhorst, 2000; Elhilali et al., 2003; Steinmetzger et al., 2019). Conventionally, such psychoacoustic experiments have been performed on a small number of listeners with known audiological characteristics, and in controlled laboratory environments. In this realm, individual variability is a nuisance parameter. In recent years, studies with a larger number of participants have exploited individual differences to understand foundational aspects of auditory processing (McDermott et al., 2010; Bharadwaj et al., 2015; Whiteford et al., 2020). However, large-N studies are difficult to perform in a laboratory environment. Web-based platforms provide an attractive alternative to quickly and cost-effectively perform studies with a large number of participants (Woods et al., 2015; Gosling and Mason, 2015). Furthermore, online platforms offer tremendous advantages in allowing researchers to access diverse and representative subject pools, perform cross-cultural research, and serve populations that may otherwise be unable to participate by visiting laboratory facilities (Henrich et al., 2010; Woods et al., 2013).

In the auditory domain, online platforms are being increasingly used to complement lab-based work (Lavan et al., 2019; McPherson and McDermott, 2018; Zhao et al., 2019). The COVID-19 pandemic has also hastened the auditory research community’s trend towards adopting online platforms. Indeed, the Psychological and Physiological Acoustics Technical Committee of the Acoustical Society of America established a Remote Testing Task Force to collect and curate information aimed at enhancing the practicality and quality of research conducted via remote testing approaches (Peng et al., 2020). Yet, to date, the literature on web-based testing of auditory perception is limited and consists primarily of examples probing higher-level cognitive aspects of auditory perception rather than low-level sensory processing. This may be in part because although studies in other domains have demonstrated that online measurements can yield high-quality data (Clifford et al., 2014; Woods et al., 2015), there is cause for skepticism about the quality of web-based psychoacoustics data for low-level aspects of auditory processing which may be more sensitive to the testing parameters and participant characteristics. Beyond the typical concerns regarding participant engagement and compliance that apply in all domains, web-based auditory experiments also have to contend with sacrificing control of sound calibration and the listening environment (e.g., ambient noise). With web-based testing, participants use their own computers and perform tasks in the environments available to them. Furthermore, given that hearing loss is highly prevalent, for example one in eight United States individuals aged 12 or older have hearing loss in both ears (Lin et al., 2011), it is likely that subject pools recruited online will consist of a significant number of individuals with hearing loss. Because hearing loss influences many aspects of auditory perception (Moore, 2007), procedures that can classify the hearing status of anonymous participants will considerably enhance the interpretability of web-based psychoacoustic measurements.

In the present study, we address these gaps by developing and validating approaches to screen anonymous participants for engagement, headphone use, and putative normal-hearing status, and test whether web-based psychoacoustic measurements can yield results comparable to lab-based procedures. While still far from the highly controlled settings in the lab, the use of headphones (or insert earphones) by participants improves the standardization of sound presentation. Headphones can attenuate ambient noise, exclude acoustic filtering that might result from the physical configuration of speakers and room reverberation, and allow for ear-specific stimulus presentation. Woods et al. (2017) developed a loudness-perception-based procedure to check for headphone use by exploiting the fact that antiphasic stimuli presented from two free-field speakers partially cancel each other, reducing the overall intensity. Here, we supplemented this procedure with a binaural hearing task that is difficult to perform with one or two free-field speakers, and difficult with monaural use of earphones, but easy with stereo headphones for individuals with normal hearing and most forms of hearing loss. We also developed and validated procedures to classify the hearing status of the participants. While conventional audiometry is infeasible without calibrated hardware, there is considerable literature on hearing screening over the telephone using suprathreshold stimuli (Smits et al., 2004; Watson et al., 2012). Inspired by this literature, we implemented participant screening by combining scores in a web-based word-recognition-in-babble task with self reports of hearing status. The choice of cutoff scores for the word recognition task were guided by a meta analysis of the literature and validation experiments. We validate the hearing-status classification procedure in a cohort of individuals with an audiological diagnosis of hearing loss. Taken together, our screening procedures help filter our subject pool for good engagement, putative normal-hearing status (if desired), and compliance with instructions to use stereo headphones. Furthermore, we validate our overall procedures by comparing the performance of screened participants on a range of classic psychoacoustic tasks with lab-based results obtained from subjects with known audiometrically normal-hearing status and prior literature.

Finally, it should be noted that the multistep screening approach used here introduces logistical challenges for anonymous participant tracking, providing a simplified experience and study flow for participants, and fair compensation that is tailored to each participant’s trajectory through the dynamically chosen components of the study. To overcome these challenges, we developed a web application using the Django framework (https://www.djangoproject.com/, Django Software Foundation) with custom features that help seamlessly integrate with the participant recruitment workflow provided by Prolific (https://www.prolific.co). In the following sections, we describe our custom infrastructure, provide further details about our overall approach, and outline the results of our validation experiments.

## Materials and Methods

### Participants for Lab-Based Testing

All lab-based human subject procedures were conducted prior to the COVID-19 pandemic, and in accordance with protocols approved by the Institutional Review Board (IRB) and Human Research Protection Program (HRPP) at Purdue University (Protocol # 1609018209). All participants had thresholds of 25 dB HL or better at audiometric frequencies in the 0.25—8 kHz range in both ears. Participants provided informed consent, and were compensated for their time performing the comodulation masking release (CMR) task. The CMR data acquired in the lab were compared to the CMR data obtained with identical stimuli from anonymous participants using web-based procedures.

### Participant Recruitment and Pre-Screening for Web-Based Testing

All web-based human subject procedures were conducted in accordance with protocols approved by the Institutional Review Board (IRB) and Human Research Protection Program (HRPP) at Purdue University (Protocol # IRB-2020-876). Participants were recruited anonymously via Prolific (https://www.prolific.co). Prolific is an online human subject marketplace where researchers can post webbased tasks (Peer et al., 2017). Anyone can sign up to be a participant and complete the tasks in exchange for payment. Individuals who sign up for participant accounts are requested to answer a series of “About You” questions. Prolific allows researchers to pre-screen participants based on their responses to different items in the “About You” section, such that only participants meeting the criteria are shown the study. Prolific also implements an approval process where researchers can “reject” participant submissions if there is clear evidence of poor engagement or poor compliance with instructions. Prolific tracks participants’ proportion of approved and rejected submissions. Prolific’s documentation provides examples of fair and unfair “attention checks” to guide the design of procedures to probe engagement. For our experiments, the pre-screening criteria required participants to:

1. be US/Canada residents and native speakers with English as first language (because we use speech stimuli spoken with North-American accents),
2. be in the 18–55 year age range,
3. have prior participation in at least 40 other studies on Prolific, and
4. have more than 90% of their previous Prolific submissions approved

For studies intended to recruit participants with normal hearing, another Prolific pre-screening criterion was that they should have answered “No” to the question “Do you have any hearing loss or hearing difficulties?”

Each participant’s Prolific ID served as the identifier for all data associated with that participant. This allowed for tracking the same individual across multiple visits to our web application. Participant compensation was a fixed amount for each task in the range of $8 to $11 per hour based on the median (across participants) time taken to complete the task.

### Infrastructure

Individual psychoacoustic tasks were created using jsPsych (https://www.jspsych.org/), a free and open-source JavaScript library for running behavioral experiments in a web browser (De Leeuw, 2015). The library is well-suited for trial-based task design and includes an array of “plugins”, each implementing a different interface for stimulus delivery and participant interaction. We used and adapted the audio plugins available in jsPsych v6.1.0.

Using jsPsych, all audio-based tasks were set up to begin with a set of instruction screens that introduced the participant to the task at hand. Following the instructions, participants landed on a volume-setting screen. This volume-setting screen instructed them to set the volume level on their computer to a low value (10-20% of the maximum on their computers). On the next screen, they were instructed to hit “play” on a calibration sound, and adjust the volume level up to a clearly audible and comfortable level while the calibration sound is playing. The calibration sound for each task was chosen to have similar spectro-temporal characteristics as the stimuli in the task (often just long strings of stimuli from the task). The stimuli for all audio trials were then presented at a level not more than 6 dB of the calibration sound. This ensured that the stimuli were clearly audible at a comfortable suprathreshold level. All audio tasks reported in this study involved stimulus presentation followed by a request for a button-click response (classic n-alternatives forced-choice trials). Feedback was provided in all audio tasks except the headphone screening using a green thumbs up for correct responses, and a red thumbs down for incorrect responses. Each audio trial was implemented using a modified version of the “audio-button-response” plugin, where the modification using javascript was done to disable (grey out) the response buttons until the audio was done playing. Because all audio tasks had a similar structure and used the same jsPsych plugins, the Django app (described below) was set up to take a JSON format (https://www.json.org/) text file with stimulus information and automatically convert it to appropriate jsPsych javascript code. This allowed for lab members who were unfamiliar with javascript to easily design their tasks.

While individual tasks were designed using jsPsych, the full-featured study design involved a combination of (1) a consent and Prolific ID confirmation page, (2) a demographics and hearing-status survey, (3) multiple individual audio tasks implemented using jsPsych with dynamic constraints on task sequence (e.g., present task #2 if subject scores more than X% in condition 1 of task #1, else conclude and debrief), (4) informational pages that showed the participants where they were within the study sequence and how

they could exit the study at any time with partial compensation, and finally (5) a debriefing page at the end that displayed the compensation amount for each subject based on their particular trajectory through the study, and redirected them to submit a “completion code” back to Prolific for study completion (or alternately request them to return the study with partial completion under certain conditions described in the Headphone-use Screening section). On the experimenter side, it was necessary to set up individual web pages to upload task information, and a page with an interface that allowed for stringing different tasks into a full study with conditional flow constraints. Pages were also set up for downloading response data, and tracking study progress and subject compensation amounts.

This complex study flow was made possible using Django (https://www.djangoproject.com/), which is a free and open-source Python-based framework for developing web applications and controlling server behavior (so-called “back-end” logic that complements our”front-end” jsPsych JavaScript). The Django application was set up to serve the consent form, the survey pages, and the individual jsPsych task pages as the subject proceeded through the study. Pages where lab members can upload the JSON files with task information and create studies were also set up. Pages other than jsPsych tasks were created using simple HTML and styled using Bootstrap (https://getbootstrap.com/). For the jsPsych tasks, the Django app was also set up to automatically and asynchronously (i.e., without refreshing the page) extract single-trial data as it was generated within jsPsych and write it to a SQLite (https://www.sqlite.org/) database on our server. The app also performed calculations (e.g., performance scores for screening tasks) to decide task flow and compensation. Each study was assigned a random alphanumeric slug as URL, and this link was posted on Prolific. Prolific allows the use of URL parameters to automatically extract the Prolific ID of each participant, which the Django app was set up to take advantage of. As a result, participants were just asked to confirm their Prolific ID rather than enter it manually.

Note that while Django is a general purpose web app framework, it offers many out-of-the-box features that made it a convenient choice to complement the capabilities of jsPsych. In particular, Django comes with functionality for working with databases using python objects (knowledge of SQL is not necessary), secure authentication capabilities that allows for logins for lab members, fine-grained permission control on creating tasks/studies and viewing results, and the ability to use sessions and cookies to track participants anonymously through various parts of the study. Importantly, the API provided by Django cleanly separates the rendering of front-end HTML/javascript from the back-end calculations and data handling, making it possible to seamlessly mix jsPsych tasks with other kinds of pages and server control. The Django app was hosted on a virtual private server (VPS) rented from Linode (https://www.linode.com/) and served by Apache2 (https://httpd.apache.org/) on Ubuntu Linux (https://ubuntu.com/). Communications between our server and subject browsers were encrypted using free SSL capabilities provided by Let’s Encrypt (https://letsencrypt.org/). A working example of our Django app, which includes a ”demo” study, can be viewed at https://www.snaplabonline.com. The source code for the application is available on GitHub (see section on Code Availability). The key resources used to build the infrastructure for web-based testing and recruit participants are illustrated in Figure 1A.

**Figure 1.**
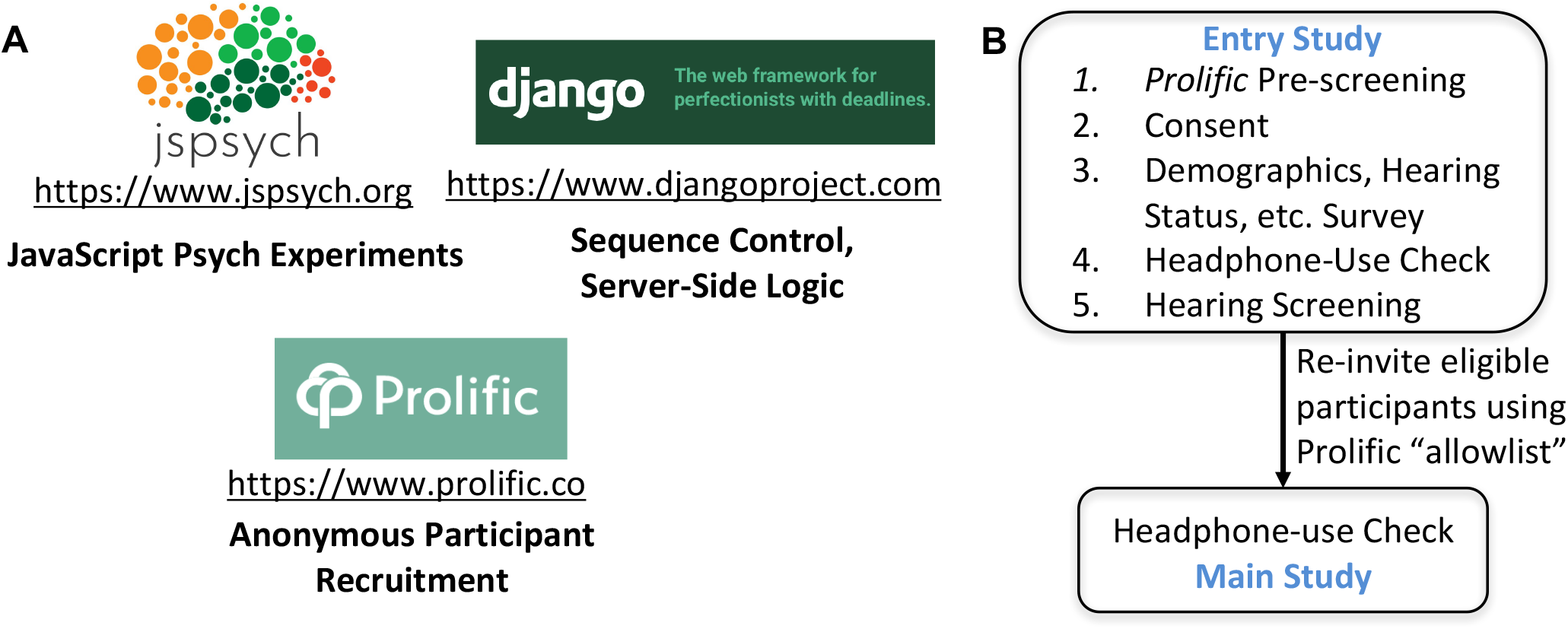
Components of the infrastructure for web-based psychoacoustics. (A) Individual tasks were created using jsPsych, an open-source javascript front-end library. Dynamic sequences of psychoacoustic tasks were implemented using Django, an open-source library for web applications, and combined with consent pages, questionnaires, and debriefing pages. Subjects were recruited via Prolific.co, a web-based human-subject marketplace. (B) A number of screening procedures were implemented to select participants for stereo headphone use, (putative) normal-hearing status, and attentive engagement via an entry study. Participants meeting criteria were re-invited to participate in the validation experiments using Prolific’s allowlist feature.

### Headphone-Use Screening

Participants were instructed to use stereo headphones in a quiet room. While compliance can generally be expected, it is possible that some participants do not comply and instead use either single-channel headphones/earphones or free-field speakers. Therefore, to objectively check for stereo headphone use, we used a combination of two tasks (six trials each). In the first task, participants were instructed to identify the softest of a sequence of three low-frequency tones. The target tone was 6 dB softer than the two foil tones, but one of the foil tones was presented with opposite phases to the left and right channels (Woods et al., 2017). Woods et al. (2017) reasoned that if a participant used a pair of free field speakers, acoustic cancellation would result in a greater attenuation of this “antiphase” foil tone compared to the target, which would cause the subject to report the wrong tone as the softest. While this is true to some extent for typical two-channel free field speakers, in general, the effectiveness of the acoustic cancellation would depend on the speaker configuration and the physical set up (e.g., distance, orientation, number of channels etc.). Crucially, if the participant used a single-channel free-field speaker or just one-channel headphone/earphone, the foil would be ineffective. To catch participants who use a single channel set up, we added a second task where participants had to report whether a low-frequency chirp (150–400 Hz) embedded in background low-frequency noise was rising, falling, or flat in pitch. The stimulus was designed such that chirp was at pi-phase between the left and right channels, whereas the noise was at zero phase (i.e., so-called “*N* 0*Sπ*” configuration). Importantly, the signal-to-noise ratio (SNR) was chosen such that the chirp would be difficult to detect with just one channel, but easily detected with binaural headphones owing to the so-called binaural masking level difference (BMLD) effect that would yield a substantial masking release (Licklider, 1948). The use of a low-frequency chirp was advantageous in two ways: (1) If a subject used two-channel free-field speakers, not only would the BMLD benefit be absent, but the same acoustic cancellation effect as described in Woods et al. (2017) would result in a further reduction in the SNR. (2) The test would be applicable even to individuals with audiometric hearing loss. This is because most individuals with hearing loss have a sloping audiogram with high-frequency loss (Parthasarathy et al., 2020), and the BMLD effect is known to be largely preserved when the low-frequency loss is not substantial (Jerger et al., 1984). Participants had to correctly respond to five out of six trials in each task to pass the headphone screening and proceed to the other tasks in the study. One exception to this was the study used to validate our hearing-screening procedures where we explicitly sought to recruit individuals with a diagnosis of audiometric hearing loss; the cut-off was relaxed to four out of six trials for that cohort. A recent study validated headphone screening procedures that similarly relied on binaural processing, and showed that combining complementary tests can boost the overall selectivity of headphone screening (Milne et al., 2020).

Note that headphone screening procedures place a logistical challenge when subjects are recruited through Prolific. Our interpretation of Prolific’s current policies (which we fully support) was that we can only exclude participants mid-way through a study without pay if there is clear evidence of non-compliance with instructions or clear evidence of inattentive engagement. However, the headphone checks of the nature used in this study cannot be expected to have 100% sensitivity and specificity. Indeed, it is possible that a small number of participants fail the headphone screening even if they comply with the instructions to use stereo headphones. This could happen if for instance, unbeknownst to the subject, their headphone (or computer settings) did not separate left and right channels adequately, or if they had asymmetric hearing loss, or if they did not understand the task well enough right from the first trial to meet threshold for passing. Thus, when a participant failed headphone screening, the Django app concluded the study and showed a debriefing page explaining that we were unable to verify headphone use (“quality check failed”) and instructed the participant to “return” the study to Prolific. Such participants were manually compensated for the time spent on the headphone screening (compensation amount tracked by the Django app) using the “bonus payment” feature on Prolific.

### Hearing Screening

It is well known that individuals with audiometric hearing loss show reduced performance in a range of suprathreshold tasks, the best known example of which is perhaps speech understanding in noise. Difficulty understanding speech in noise is the most common audiological complaint among individuals who are eventually diagnosed with hearing loss (Blackwell et al., 2014). Thus, while conventional threshold audiometry is infeasible without calibrated hardware, such suprathreshold tasks may plausibly be used to screen for hearing loss. Indeed, identification of spoken digits in background noise has been validated for hearing screening via telephone in many countries by comparing the threshold SNR for digit identification to audiometric thresholds (Smits et al., 2004; Watson et al., 2012). Along the same lines, we developed hearing screening procedures using material from the modified rhyme test/MRT (House et al., 1963) presented in 4-talker babble. The MRT materials include 50 lists of monosyllabic words, where each list consists of six monosyllabic word alternatives that differ in just the first or the last phoneme. The MRT is thus convenient for administration with a web-based interface by virtue of allowing for a 6-alternatives forced choice response. Despite the closed-set nature of the response, the MRT is known to capture the suprathreshold deficits resulting from hearing loss (Elkins, 1971; Bell et al., 1972; Brungart et al., 2014). To use the MRT-in-babble as a hearing screener, suitable cutoff scores must be determined below which an individual would be considered as failing the screening. Any given performance cutoff represents a tradeoff between sensitivity and specificity for detecting hearing loss, with the precise trade-off relationship dependent on the effect size of hearing loss on speech-in-noise scores. To determine a suitable performance cutoff, we conducted a meta analysis of 15 studies that reported speech-in-noise scores for various materials across normal-hearing listeners and individuals with hearing loss (pure tone average thresholds > 30 dB HL). Inverse variance pooling was used to estimate the effect size using the bias-corrected procedure outlined in Hedges (1982). The efficacy of the hearing screening procedure was then validated by measuring the pass rates among individuals with a history of hearing loss as diagnosed by an audiologist or otologist.

### Psychoacoustic Tasks

The primary experimental strategy used to investigate the validity and viability of web-based psychoacoustics was to compare results from web-based administration of a range of classic psychoacoustic tasks to corresponding data from the same (or substantially similar) tasks in controlled lab-based settings. Specifically, we used the following measures to validate our web-based procedures:

1. Fundamental-frequency (F0) discrimination (F0 difference limens; F0DLs)
2. Gap detection (duration thresholds)
3. Interaural time difference (ITD) threshold sensitivity
4. Interaural level difference (ILD) threshold sensitivity
5. Comodulation masking release (CMR) effect
6. Effect of SNR on word identification, and
7. Consonant confusion patterns in background noise

The tasks employed target a range of phenomena pertaining to sensory processing that have been classically studied in the psychoacoustics literature. The tasks included detection of subtle diotic and dichotic cues (F0DLs, Gap/ITD/ILD thresholds), the use of modulation cues for auditory scene analysis (CMR), and speech categorization with no syntactic or semantic context (monosyllabic word identification and consonant confusions). The specific stimuli employed for each task are described alongside the measured performance trends in the Validation Experiments and Results section.

### Overall study Sequence

To integrate our screening procedures and psychoacoustic tasks with the workflow facilitated by Prolific, the data collection procedures were mapped to two separate studies on Prolific (Figure 1B). In addition to the pre-screening based on Prolific’s “About You” bank of questions, Prolific also allows for custom pre-screening where researchers can upload an “allowlist” of Prolific IDs to restrict which participants are shown the study. This allowlist feature enables longitudinal designs where researchers can store the IDs of participants from one study, apply any custom criteria to the data from that first study, and re-invite participants who meet criteria for follow-up studies. We leveraged this feature to implement our hearing screening procedures within an entry study, and the participants who passed the screening were re-invited to participate in studies that contained the other main psychoacoustic tasks (Figure 1B). Headphone screening procedures were included in both the entry study and the follow-up studies.

## Validation Experiments and Results

### Validation of hearing screening procedure

For hearing screening, we applied cutoffs to percent-correct scores obtained for recognition of words from the modified rhyme test (MRT; House, 1963). The words were presented with the carrier phrase “Please select the word “ in 4-talker babble with matching long-term average spectrum at four different SNRs (10, 5, 0, and *-*5 dB). To help choose appropriate cutoffs, a meta-analysis of the literature was conducted. This helped obtain an initial estimate of effect-size with which speech-in-noise tests generally separate listeners with normal hearing (NH; pure-tone average/PTA thresholds ≥25 dB) from those with hearing loss (HL; PTA ≥ 30 dB). 15 studies were selected for inclusion based on the following criteria:

1. Use of English-language materials
2. Reporting of either percent-correct speech identification for a fixed SNR away from floor and ceiling, or the SNR at which a fixed percent-correct was obtained
3. Inclusion of NH and HL groups within the same study
4. Explicit reporting of mean and standard deviations in each group
5. Use of one or more of the following standard speech-testing materials: Bamford-Kowal-Bench Speech-in-Noise Test or BKB-SIN (Niquette et al., 2003), the Quick Speech-in-Noise Test or QuickSIN (Killion et al., 2004), Words-in-Noise test or WIN (Wilson, 2003), Hearing in Noise Test or HINT (Nils-son et al., 1994), NU-6 words in noise (Auditec Inc.), or the MRT

Although the meta analysis was not exhaustive, and the speech-testing material was non-uniform across the studies chosen, we considered the analysis adequate to provide an initial estimate of cutoffs for our purposes. This is because the actual selectivity of our web-based hearing screening was separately quantified by applying the chosen cutoffs to data from a cohort of individuals with a diagnosis of hearing loss. The number of studies (i.e., 15) included in the analysis was also considered adequate because the effect size was estimable with a narrow confidence interval (Figure 2A) using bias-corrected inverse variance pooling procedures (Hedges, 1982). The meta-analysis revealed a large effect size of g = 2.05 based on the pooled-variance estimate (Figure 2A), and that the standard deviation of scores in the HL group was a factor of three larger than the standard deviation in the NH group. Using this standard deviation ratio and overall effect-size estimate, it is possible to theoretically calculate the receiver-operator characteristics (ROC curves) that can be expected when using a cutoff procedure to blindly classify the hearing status of an individual participant. This theoretical ROC curve is shown in Figure 2B. These initial estimates suggested that if we choose a cutoff such that we exclude 20–40% of NH participants, we would be able to exclude around 90% of participants with hearing loss. Because our initial pool in web-based testing would likely include some participants with hearing loss, and because it is possible that web-based procedures yield larger across-participant variance, we chose our initial cutoff conservatively to exclude 35% (i.e., closer to 40% rather than 20%) of the entry pool. Accordingly, our hearing screening procedure deemed participants who scored 100% at 10 dB SNR, *≥* 83% at 5 dB SNR, and *≥* 75% at 0 dB SNR to have “normal” hearing. Note that setting performance cutoffs on a task also inherently filters for participants showing good compliance with instructions and focused attention. To further filter for focused engagement, we also eliminated any participants who registered more than two “blur” events during the course of this 6–7 minute long MRT test. This was possible because jsPsych automatically records a few different kinds of user interaction events. A “blur” event occurs when the user clicks on another window or tab during the experiment, indicating that they are no longer interacting with the experiment.

**Figure 2.**
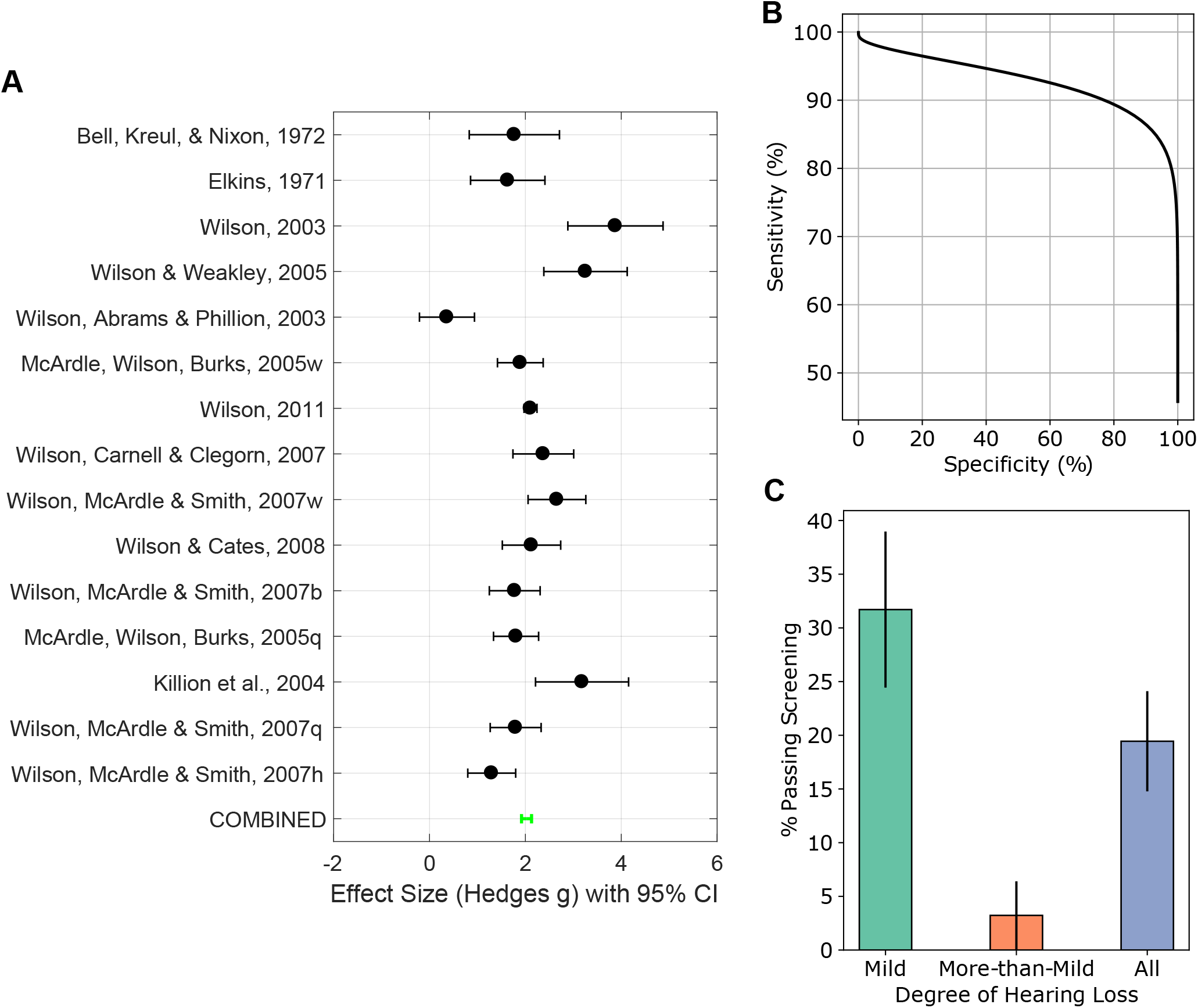
Validation of the hearing screening procedure. (A) A meta-analyis of 15 speech-in-noise studies in the literature suggested that such tasks yield a large effect size separating individuals with normal hearing (NH) and hearing loss (HL). (B) Theoretical calculations based on results from panel (A) predict the expected tradeoff between sensitivity and specificity for the detection of hearing loss. This was used to determine cutoff scores for the hearing screening procedure. (C) Actual selectivity of our hearing screening procedure was quantified in a cohort of subjects with a professional (clinical) diagnosis of hearing loss. Sensitivity for any degree of HL was about 81%. Sensitvitiy for more-than-mild HL (i.e., moderate, moderately-severe, or severe) was > 95%. Error bars in panel (C) indicate standard deviation of estimated pass rates from *N* = 72 subjects.

To estimate the actual selectivity of our hearing screening procedure, we applied the procedure to a separate pool of participants who met all of the following conditions: They (1) self-identified as having hearing loss or hearing difficulties on Prolific, (2) were subsequently able to confirm through our survey that they had indeed received a diagnosis of hearing loss from a medical professional (audiologist or otologist), and (3) were able report their diagnosed degree of HL with high confidence. Of this pool of *N* = 72 participants, only 19% exceeded our cutoff (i.e., the overall sensitivity for detecting any degree of HL was about 81%). An examination of the degree of HL among those who passed our hearing screening revealed that the majority of hearing-impaired participants that our test failed to catch only had a mild HL. Specifically, while 31% of subjects reporting a clinical diagnosis mild HL were able to meet cutoff, only 3% those with more-than-mild HL (i.e., clinical diagnosis moderate, moderately severe, or severe HL combined) were able to meet cutoff (Figure 2C). Taken together our results confirm that we obtain > 65% specificity (by choice of cutoff, less than 35% of NH individuals will fail hearing screening), > 80% sensitivity for correctly excluding subjects with any degree of HL, and > 95% sensitivity for correctly excluding subjects with more-than-mild HL. Thus, although by no means a substitute for audiometric screening, our suprathreshold screening based on MRT scores is quite successful in filtering the subject pool for near-normal hearing.

To improve our hearing screening even further, we supplemented this MRT-based screening with the requirement that participants had to explicitly deny having hearing loss in our demographics survey. We also required that participants had to deny having any neurological disorders or persistent tinnitus. A Prolific-style “allowlist” of entry-study participants who met all of our criteria was maintained, and the “main” studies were only open to individuals on this list (Figure 1B). Next, we measured the performance of re-invited participants on a range of classic psychoacoustic tasks to test whether web-based psychoacoustics yields results comparable to lab-based data.

### Absolute sensitivity measurements using diotic stimuli

The first series of measurements probed absolute psychoacoustic sensitivity to subtle fundamental frequency shifts (perceived as pitch variations) and gap duration cues for *N* = 100 participants in our allowlist. Fundamental frequency (F0) discrimination was tested for a harmonic complex tone with an F0 of 110 Hz, using stimulus parameters resembling a recent large-scale lab-based study (Madsen et al., 2017). Specifically, F0 difference limens were measured using a three-interval 3-alternative forced-choice (3AFC) paradigm. Each interval consisted of a sequence of four 200 ms-long complex tones ramped on and off using 20 ms-long Hann tapers. The first and third tone in the target interval had a symmetric F0 shift centered at 110 Hz. All four tones in the two foil intervals had a constant F0 of 110 Hz. F0 difference sensitivity was measured for each participant using the method of constant stimulus for various values of F0 shift from 0.05 Hz to 3.2 Hz in geometric steps. Psychometric functions were fitted using a bayesian approach with a beta-binomial model (Schütt et al., 2016). Numerical integration procedures implemented in Psignifit 4 were used to estimate threshold, slope, and lapse-rate parameters using the default priors in the software, and with chance level set to 33% (Schütt et al., 2016). The average and 95% (posterior) confidence interval for the F0 difference limen, defined as the 70% point on psychometric function, were estimated (Figure 3A). The F0-difference limen was about 0.4%, showing that the exquisite F0 sensitivity of typical normal-hearing human subjects was estimable using web-based testing. These results are a close match to lab-based estimates (Madsen et al., 2017).

**Figure 3.**
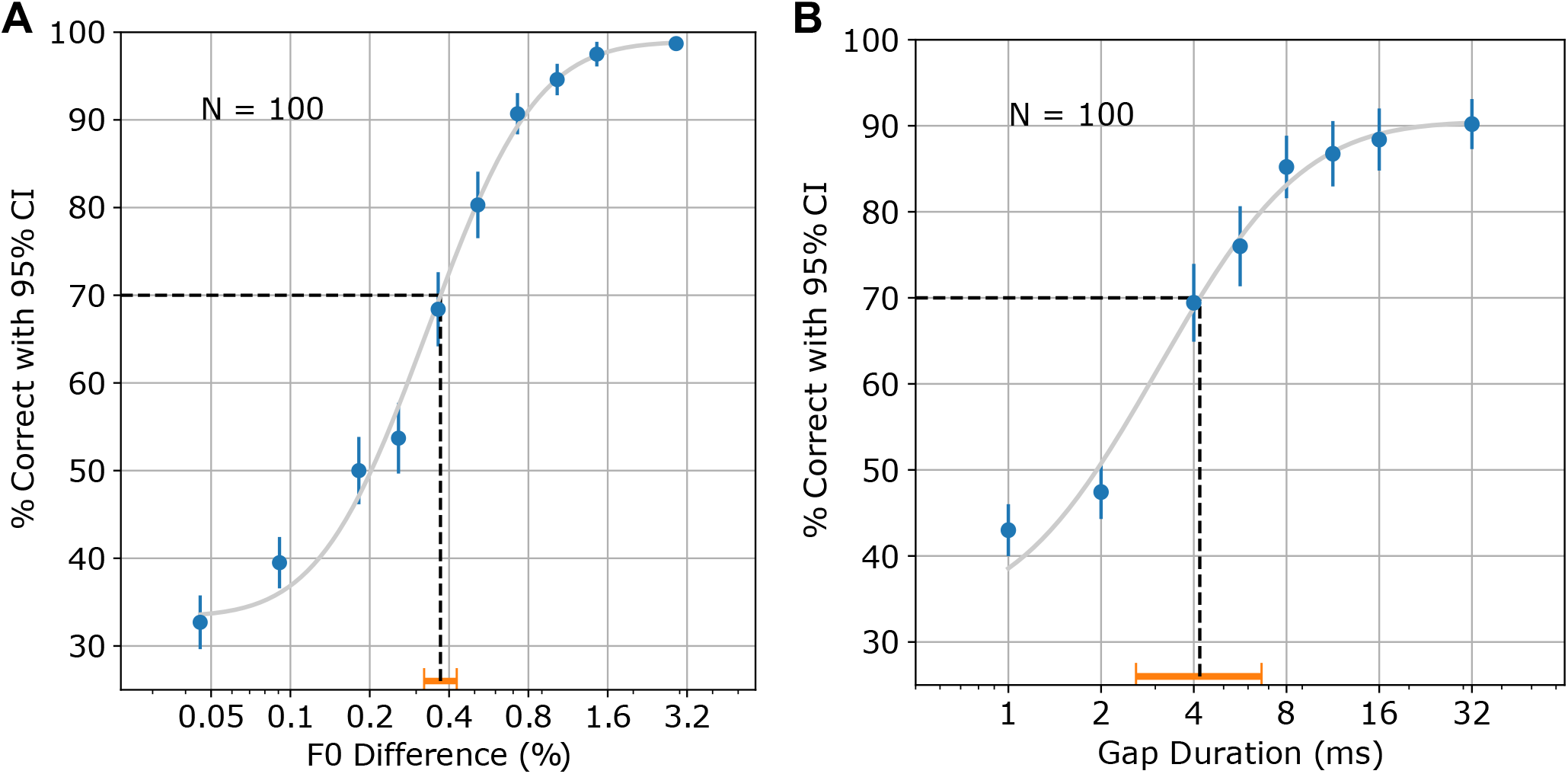
Validation of web-based testing for classic psychoacoustic tasks with diotic stimuli. Psychometric functions were measured for fundamental-frequency (F0) discrimination (A), and gap detection (B). F0 discmination limens were around 0.4% for F0 = 110 Hz, and gap thresholds were around 4 ms, consistent with lab-based data with similar stimuli. Horizontal error bars in orange indicate the 95% confidence interval of the mean threshold estimated from *N* = 100 subjects.

Gap-duration sensitivity was measured using a simple 3AFC detection task for short gaps in 4 kHz tones embedded in background noise. The stimulus parameters were chosen to resemble a recent large-scale lab-based study (Patro et al., 2020). The background noise ranged from 0.5 octaves below to 0.5 octaves above 4 kHz and was 10 dB below the 4 kHz tone in intensity (i.e., SNR = 10 dB). The tonal segments before and after the short gap were each 125-ms long and were ramped on and off using 1-ms-long Hann tapers. The noise itself started 50 ms before the first tone segment and was gated on with a 10 ms Hann taper. The gap duration in the target interval was varied, whereas the two foil intervals always had a gap duration of 0 ms. Detection accuracy was measured for target gap durations ranging from 1 to 32 ms in geometric steps. Once again, psychometric functions were fitted using the bayesian approach. Results demonstrated exquisite sensitivity to gaps with mean thresholds of about 4 ms (Figure 3B), which are well in line with lab-based results from Patro et al. (2020).

Taken together, our results suggest that despite using participants’ own hardware in their own listening environments, absolute psychoacoustic sensitivity measures matched highly controlled lab-based data for diotic stimuli.

### Absolute sensitivity to interaural differences

Next we asked whether web-based measurements can replicate lab-based data when ear-specific stimuli need to be delivered. To this end, we probed absolute sensitivity to interaural cues. Humans are known to be sensitive to interaural time differences in low-frequency sounds that are an order of magnitude or more smaller than the width of typical neural spikes (about 1 ms), and sensitive to interaural level differences that occur in the environment for a source that is displaced from the azimuthal center by as few as a couple of degrees for high-frequency sounds (Moore, 2012). We sought to test whether such sensitivity can be robustly demonstrated with web-based testing. Interaural time difference (ITD) sensitivity for 500 Hz tones was assessed using a two-interval task where the leading ear was switched between the intervals. Participants were asked to report whether the sound jumped from left-to-right or right-to-left in a 2-alternative forced choice (2AFC) design. Accuracy was measured for ITD values ranging from 2 microseconds to 128 microseconds in geometric steps. Psychometric curves were fitted using the bayesian approach as before for data collected from *N* = 200 subjects who had passed our screening procedures. ITD thresholds (defined as 75% correct points) in the vicinity of 25 microseconds were obtained (Figure 4A), consistent with classic literature (Zwislocki and Feldman, 1956; Klumpp and

**Figure 4.**
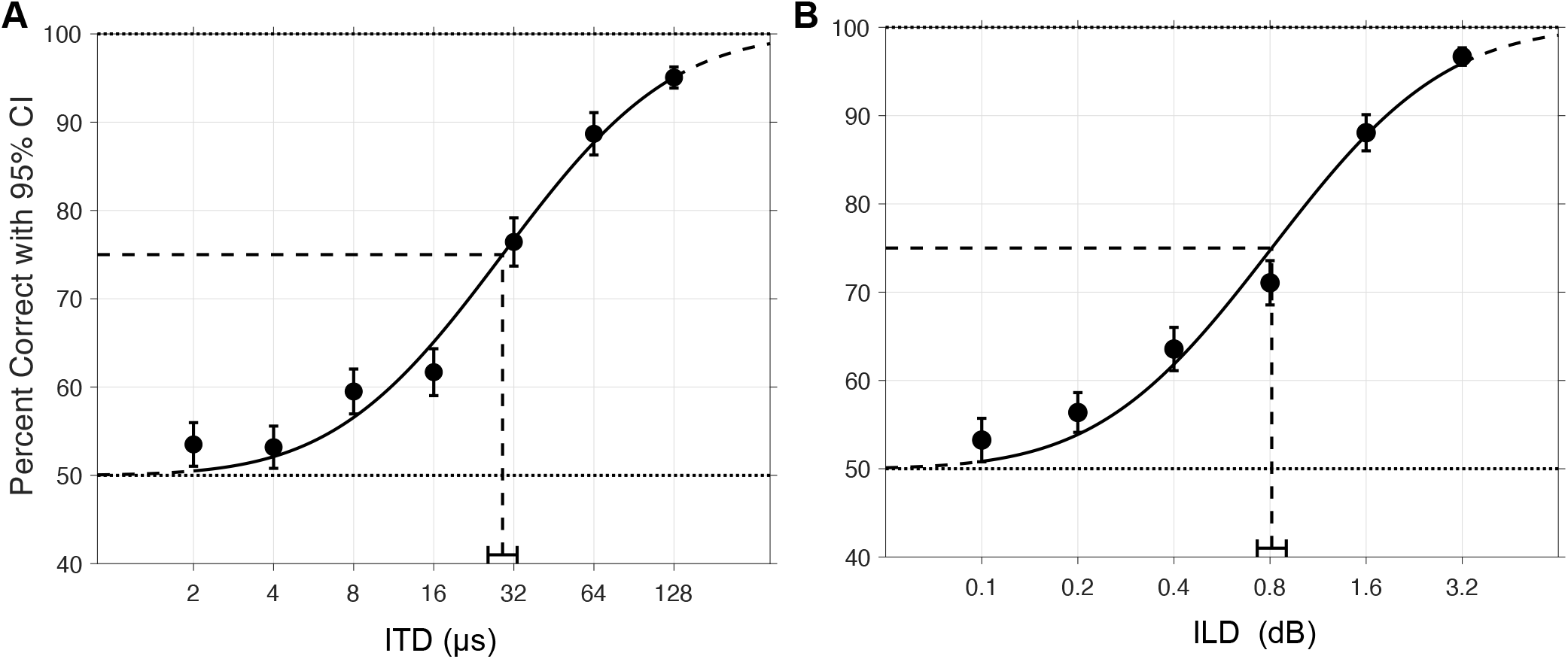
Validation of web-based testing for classic interaural difference detection tasks. Psychometric functions were measured for interaural time-delay (ITD) detection (A), and interaural level-difference (ILD) detection (B). ITD thresholds were around 25*µ*s for 500 Hz tone bursts, and ILD thresholds were around 0.8 dB for 4 kHz tone bursts, consistent with classic literature. Horizontal error bars indicate the 95% confidence interval of the mean threshold estimated from *N* = 200 subjects.

Along the same lines, interaural level difference (ILD) sensitivity for 4 kHz tones was assessed using a two-interval task, where the ear with the more intense tone was switched between the intervals. As with the ITD task, participants were asked to report whether the sound jumped from left-to-right or right-to-left (2AFC). Data from *N* = 200 subjects revealed that ILD thresholds were around 0.8 dB, consistent with classic literature (Mills, 1960). The psychometric data for ITD and ILD detection are shown in Figure 4 and confirm the validity of web-based testing for probing sensitivity to binaural cues.

### Measuring scene analysis using comodulation masking release

Humans use statistical regularities in sounds to perceptually organize sound information that arrives simultaneously from multiple sound sources in the environment (Bregman, 1990). One acoustic cue that promotes the grouping of different components of a sound source into a single perceptual object is co-modulation (Darwin, 1997). Indeed, it is thought that masker sounds that contain acoustic components that are temporally coherent with a target source will perceptually interfere with target perception (Shamma et al., 2011; Viswanathan et al., 2021). The phenomenon of comodulation masking release (CMR) using simple tone-in-noise stimuli is thought to capture aspects of such co-modulation based scene analysis. For tones masked by on-frequency noise, the addition of flanking bands of noise, or a widening of the bandwidth of the on-frequency masking noise significantly improves the detection thresholds for the target tones, but only if the different masker frequency components are co-modulated (Schooneveldt and Moore, 1989). Qualitatively, co-modulated masker components are perceived as a single broadband object thereby allowing for the target (narrowband) tone to stand out. Here, we tested whether the CMR effect can be illustrated using web-based psychoacoustic measurements.

Detection thresholds were measured for 4 kHz tones embedded in narrowband 1-ERB-wide (Glasberg and Moore, 1990) 10-Hz modulated noise, with various configurations of flanking noise bands (1 ERB bandwidth, 2 ERB gap between bands; Figure 5A). CMR was calculated as the difference in the detection threshold across the different flanker conditions. In-lab (pre-COVID-19) measurements using an adaptive procedure on *N* = 40 subjects with normal audiograms (and similar age as our Prolific participants) suggested that CMR was > 3 dB for CORR - REF conditions, and > 12 dB for CORR - ACORR conditions. The psychometric functions obtained with identical stimuli using web-based measurements are shown in Figure 5B (*N* = 203). The CMR measured via web-based testing was abouot 4 dB for CORR - REF conditions, and about 17 dB for CORR - ACORR conditions, establishing that robust CMR effects are demonstrable with web-based testing as in lab-based tests.

**Figure 5.**
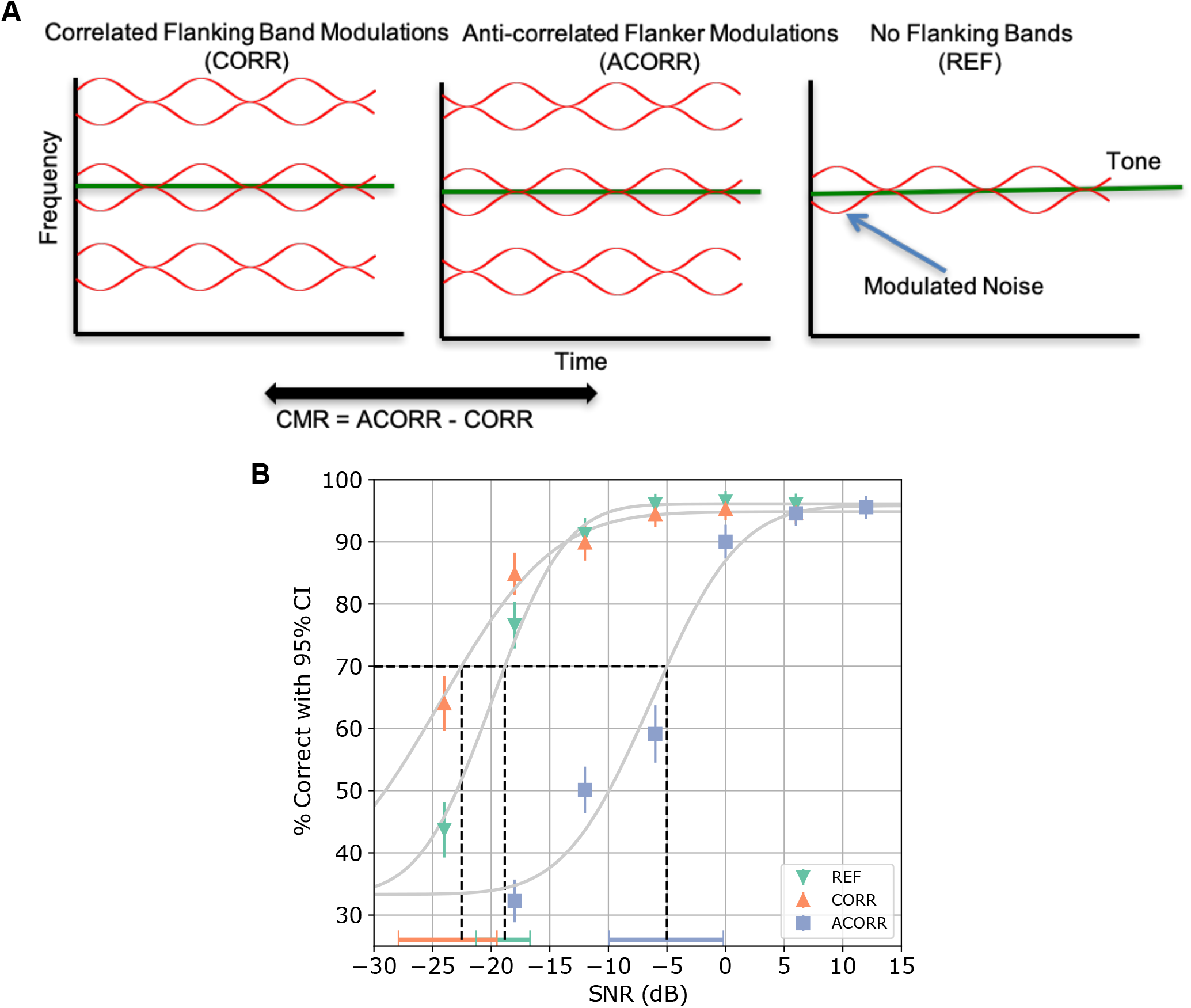
Validity of web-based testing for demonstrating the co-modulation masking release (CMR) effect. (A) CMR was measured by assessing the SNR thresholds for detecting 4 kHz tones in modulated 1-ERB-wide on-band noise for varying configurations of 1-ERB-wide noise flankers (2-ERB gap between on-band noise and flankers). The flankers were either absent (REF), modulated in a correlated manner with the on-band noise (CORR), or anticorrelated manner with the on-band noise (ACORR). The change in tone thresholds across conditions is quantified as the CMR effect. (B) Psychometric curves for tone detection in different CMR conditions was measured from N=203 subjects, yielding a clear separation between conditions. CMR for CORR-REF was about 4 dB, whereas CMR for CORR - ACORR was about 17 dB consistent with lab-based measurements with identical stimuli. Horizontal errorbars indicate 95% confidence intervals for the tone threshold in each condition.

### Word-recognition in noise and consonant identification

Finally, we sought to compare word recognition performance and consonant categorization patterns measured using web-based testing with corresponding lab-based data from the literature. Figure 6A shows the word-recognition scores in background babble from the MRT test that was used for hearing screening. Overall performance levels and trends with SNR for participants who passed our web-based hearing-screening replicate lab-based results obtained from individuals with normal audiometric thresholds (Miner and Danhauer, 1976).

**Figure 6.**
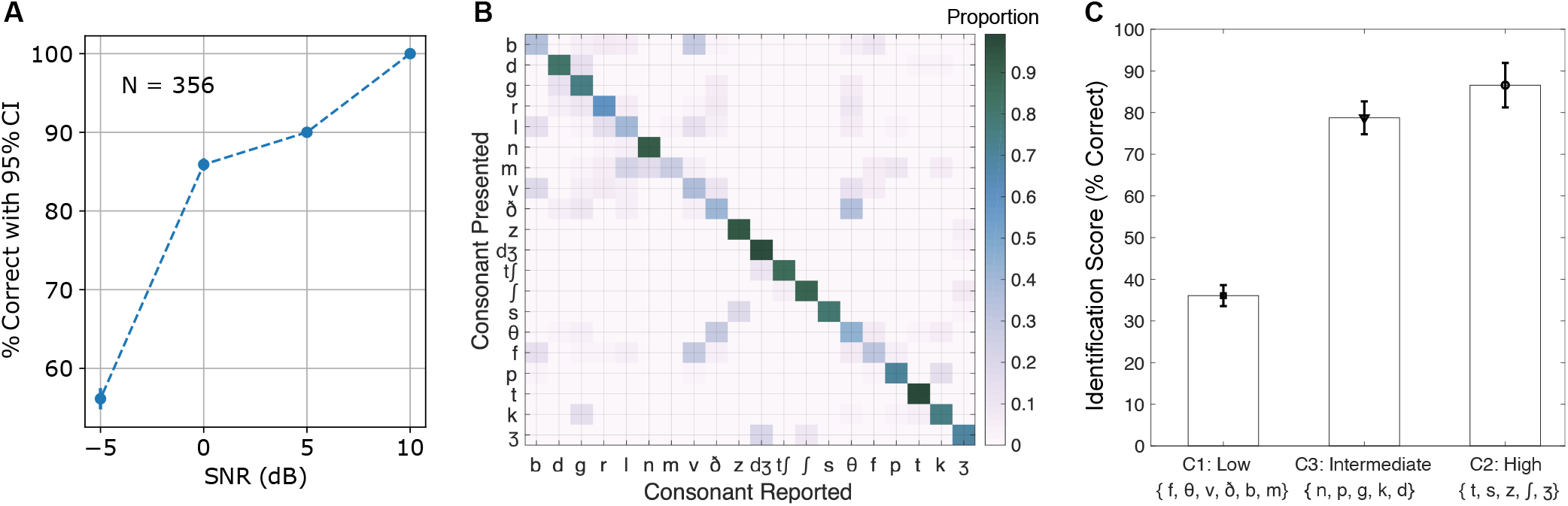
Validation of web-based testing using word-recognition scores and consonant confusion patterns. (A) Recognition scores for monosyllabic words embedded in spectrally matched (in the long-term average sense) 4-talker babble showed overall performance and SNR dependence consistent with literature (Miner and Danhauer, 1976). (B) Consonant confusion matrix for consonants presented in C/a/ context in speech-shaped stationary noise at –8 dB SNR from *N* = 295 samples across four talker voices. Confusion clusters extracted from this matrix match lab-based results (Phatak and Allen, 2007). (C) A summarization of diagonal entries from panel (B) revealed that recognition scores were not uniform across consonants. Differences across sets of consonants with low, intermediate and high scores match lab-based data (Phatak and Allen, 2007).

To probe consonant categorization and confusions (Miller and Nicely, 1955), participants were asked to identify consonants (i.e., C/a/ syllables) from the STeVI corpus (Sensimetrics Corporation, Gloucester, MA) presented in speech-spectrum stationary noise using the carrier phrase “You will mark ― please”. An SNR of –8 dB was chosen, which yielded an overall intelligibility of 66% (and hence a sufficient number of confusions for analysis). Consonant confusions from N=295 samples across four different talker voices are shown in Figure 6B. This data was compared to controlled lab-based results using similar stimuli (Phatak and Allen, 2007). Phatak and Allen (2007) found that for a fixed overall intelligibility, recognition scores varied across different consonants. Specifically, they identified three groups of consonants that they referred to as “C1”, “C2”, and “C3”, with low, high, and intermediate scores, respectively. Our results replicate this trend for the specific consonant groups that they identified, and closely match the relative values of the scores across the groups (Figure 6C). Based on a graphical analysis of confusion patterns, Phatak and Allen (2007) also identified confusion sets (i.e., sets of consonants where any consonant in a particular set is most confused with another consonant in the same set). Our results from the online experiment replicate their lab-based results in that when we used probability of confusion between a pair of consonants (from Figure 6B) as the distance metric to perform hierarchical agglomerative clustering, we obtained the same confusion sets. Specifically, when we thresholded our agglomerate to yield the same number of clusters as reported in Phatak and Allen (2007), the clusters from our data matched their report: {f, v, b, θ, ð}, {s, z}, {⎰, 3}, and {m, n}.

## Discussion

Although web-based experiments are used in many domains of behavioral research, they have not received much adoption in psychoacoustics research. This is understandable given the challenges stemming from lack of control of stimulus hardware or listening environment, and from the inability to know the hearing status of anonymous participants. Psychoacoustic phenomena pertaining to low-level sensory processing of sounds is generally thought to be sensitive to such testing parameters. Here, we developed and implemented a range of strategies to mitigate these challenges and tested whether web-based psychoacoustics is in fact viable.

For hearing screening, we used a suprathreshold word-recognition-in-babble task and documented evidence for selectivity for near-normal hearing status. Specifically, we found that by setting specificity to around 65% (i.e., exclude about 35% of all participants), we can detect hearing loss of any degree with > 80% sensitivity, and detect more-than-mild hearing loss with > 95% sensitivity. We supplemented this hearing screening with procedures to check for stereo headphone use, focused engagement (i.e., not clicking on other tabs/windows during the experiment), screening for demographics characteristics (age and English language fluency status), and self-report of normal hearing (including no persistent tinnitus) and neurological status. We then evaluated web-based psychoacoustics by comparing data from web-screened participants to lab-based data from individuals with known audiometric status, and found excellent agreement.

We found that with web-based testing, absolute sensitivity to diotic and binaural cues were robustly estimable, the main effects of changes in modulation statistics of the auditory scene were readily apparent, and that speech recognition and consonant category confusion patterns closely matched lab-based results. In particular, F0 difference limens for web-based data were on the order of 0.4%, gap detection thresholds in tonal markers was about 4 ms, ITD thresholds for low-frequency tones were about 25 microseconds, ILD thresholds for high-frequency tones was about 0.8 dB, and consonant confusions were in agreement with highly controlled studies.

Taken together, our results show that web-based psychoacoustics is viable and an excellent complement to lab-based work. Our screening procedures and psychoacoustic tasks are implemented using open-source tools (jsPsych and Django), which provide capabilities for trial-based experiment design with complex study flow. We make the source code for our custom web app available via GitHub for interested investigators to adapt to their needs. We also make a working demo of our infrastructure available.

Remote psychoacoustic testing can help overcome many challenges with lab-based testing. One important example of a challenge that remote testing can help mitigate is the restrictions on in-person experiments during the COVID-19 pandemic. Other key advantages of remote testing include the ability to collect large datasets at lower costs, the ability to include diverse and representative cohorts of subjects, to reach and serve certain groups who may otherwise find themselves excluded from lab-based research in the convenience of their homes, and the ability to do longitudinal training or monitoring over extended periods. While a range of approaches can be adopted within the remote-testing umbrella (Peng et al., 2020), the present study focused on web-based testing on anonymous participants using their own devices and web browsers in whatever listening environments were available to them. Other remote-testing possibilities that allow for greater hardware control include shipping lab-calibrated hardware to remote participants such as tablets/smartphones and a headphone (Lelo de Larrea-Mancera et al., 2020), or having participants install apps on specific consumer-grade devices with known average acoustic characteristics. We chose to focus on browser-based testing here because it requires minimal co-operation from subjects for installing specialized software and is easily scalable to a range of devices and operating systems. Although browser-based testing is perhaps the least controlled of remote testing options, our results using carefully designed procedures show that excellent outcomes can be obtained with such web-based testing. One potential limitation of our study is that our validation experiments did not directly assess the viability of web-based testing for between-subject comparisons. However, the fact that our hearing screening procedures can separate groups with normal hearing and hearing loss is encouraging in this respect.

## Declarations

### Funding

This work was supported by National Institutes of Health (NIH) grants R01DC015989 (HMB), F31DC017381 (VV), and T32DC016853 (RS).

### Author Contributions

HMB conceived the project, designed the protocols, and implemented the infrastructure. BAM performed meta analysis to guide hearing screening procedures. VV, AB, RS, and HK provided design input and contributed validation data. All authors contributed to manuscript writing.

### Code Availability Statement

Our infrastructure was developed using open-source toolboxes (jsPsych and Django). The code for our Django application is publicly available on GitHub (https://github.com/haribharadwaj/SNAPlabonline) and archived using Zenodo (Bharadwaj, 2021). A working demo of the app is hosted at https://snaplabonline.com. The license for the code is highly permissive and interested investigators are welcome to adapt as needed for their purposes. The audio files used to create the MRT stimuli were obtained from the National Institute of Standards and Technology PSCR Audio Source Files repository (https://www.nist.gov/ctl/pscr/pscr-audio-source-files). The speech materials used for the consonant confusion task were from the commercially available STeVI corpus (Sensimetrics Corporation, Gloucester, MA).

### Data Availability Statement

Data to reproduce all figures will be made publicly on Figshare (https://figshare.com/) upon peerreviewed publication.

### Competing Interests

The authors declare no competing financial or non-financial interests.

### Ethics approval

All human subject procedures were conducted in accordance with protocols reviewed and approved by the Purdue University IRB.

### Consent to participate

All subjects provided affirmative informed consent.

### Consent for publication

Human subjects provided consent to publish their de-identified data.

## Notes

### Competing Interest Statement

The authors have declared no competing interest.

